# The Oscillatory Effects of Rhythmic Median Nerve Stimulation

**DOI:** 10.1101/2020.10.23.348268

**Authors:** Mairi S. Houlgreave, Barbara Morera Maiquez, Matthew J. Brookes, Stephen R. Jackson

## Abstract

Entrainment of brain oscillations can be achieved using rhythmic non-invasive brain stimulation, and stimulation of the motor cortex at a frequency associated with sensorimotor inhibition can impair motor responses. Despite the potential for therapeutic application, these techniques do not lend themselves to use outside of a clinical setting. Here, the aim was to investigate whether rhythmic median nerve stimulation (MNS) could be used to entrain oscillations related to sensorimotor inhibition. MEG data were recorded from 20 participants during 400 trials, where for each trial 10 pulses of MNS were delivered either rhythmically or arrhythmically at 12 or 20Hz. Our results demonstrate a frequency specific increase in relative amplitude in the contralateral somatosensory cortex during rhythmic but not arrhythmic stimulation. This was coupled with an increase in inter-trial phase coherence at the same frequency, suggesting that the oscillations synchronised with the pulses of MNS. While the results show that 20Hz rhythmic peripheral nerve stimulation can produce entrainment, the response to 12Hz stimulation was largely due to the presence of rhythmic sensory evoked potentials. Regardless, rhythmic MNS resulted in synchronous firing of neuronal populations within the contralateral somatosensory cortex meaning these neurons were ‘occupied’ in processing of the afferent input. Therefore, MNS could prove therapeutically useful in disorders associated with hyperexcitability within the sensorimotor cortices.

## 1. Introduction

Neural oscillations are rhythmic variations in electrical activity which arise from the summation of synchronous postsynaptic potentials. Oscillations are categorised based on their frequency, with the alpha/mu-alpha (8-12Hz) and beta (13-30Hz) frequency bands being of particular importance when researching sensorimotor systems. In the contralateral sensorimotor cortex, oscillations in the 8-30Hz range are suppressed during movement and movement preparation, but following movement there is a beta rebound, meaning their amplitude is briefly higher than at rest (Jurkiewicz et al., 2006; Pfurtscheller et al., 1996). Corticospinal excitability is known to be increased when sensorimotor oscillations are suppressed and reduced during the post-movement beta rebound (Chen et al., 1998). Higher beta activity has also been associated with a slowing of newly initiated movements (Gilbertson et al., 2005). Therefore, beta synchrony is frequently thought of as a mechanism which promotes maintenance of the current motor set (Engel and Fries, 2010). However, a newer theory suggests that beta is a marker of motor readiness, whereby a low level of beta activity indicates a higher likelihood that a movement will be generated (Jenkinson and Brown, 2011). On the other hand, alpha (mu-alpha in the sensorimotor cortex) synchrony has been linked to the inhibition of task irrelevant areas (Brinkman et al., 2016; Jensen and Mazaheri, 2010), with the network of regions involved in a task showing a desynchronisation of alpha oscillations (Haegens et al., 2010). Various non-invasive brain stimulation (NIBS) techniques have been shown to modulate these oscillatory rhythms as well as behaviour (Pogosyan et al., 2009; Thut et al., 2011b). As a result, there is widespread interest in the use of these techniques as potential forms of therapy in a multitude of disorders.

An interesting avenue for therapy using NIBS is entrainment; the process though which neuronal assemblies become synchronised to a rhythmic stimulus train (Thut et al., 2011a). Entrainment of oscillations in the 8-30Hz range could prove therapeutically beneficial in patients with disorders characterised by sensorimotor hyperexcitability (Morera Maiquez et al., 2020b). Therefore, it is of interest that repetitive transcranial alternating current stimulation (tACS) at a frequency associated with periods of decreased corticospinal excitability can lead to a reduction in the velocity of movement (Pogosyan et al., 2009). In 2011, Thut and colleagues showed that rhythmic transcranial magnetic stimulation (TMS) of a parietal site at an individual’s preferred alpha frequency caused region specific entrainment of brain oscillations (Thut et al., 2011b). These findings were specific to rhythmic TMS and were not seen with arrhythmic or sham stimulation control conditions. Given that tACS can have similar oscillatory effects it is interesting that application of a topical anaesthetic can significantly reduce entrainment, suggesting that the effects of tACS may be due to stimulation of the somatosensory cortex via peripheral nerves rather than direct stimulation of the cortex itself (Asamoah et al., 2019). This indicates that there is potential for the same effect to be replicated through rhythmic stimulation of peripheral nerves rather than the cortex. In fact, rhythmic median nerve stimulation (MNS) at 12Hz and 19Hz (preprint) has shown similar entrainment effects in healthy participants, using sensor level EEG analysis (Morera Maiquez et al., 2020b, 2020a (preprint)). Furthermore when compared with no stimulation, delivery of 10Hz rhythmic MNS has been shown to reduce the frequency of tics in Tourette’s syndrome (TS) patients, a disorder associated with sensorimotor hyperexcitability (Morera Maiquez et al., 2020b). These are important findings as, unlike tACS and TMS, MNS is portable, cheaper and requires little training, making it an ideal technique to be used therapeutically outside of the clinic.

When a sensory stimulus (such as MNS) is delivered a transient, phase-locked electrical potential can be recorded over the sensorimotor cortex which is known as a sensory evoked potential (SEP) (Vialatte et al., 2010). When longer trains of rhythmic stimuli are delivered, we can record steady-state evoked potentials (SSEPs) which appear as a sustained response at the frequency of stimulation (Regan,1989, as cited in (Vialatte et al., 2010))(Regan, 1966). SSEPs are thought to either arise due to entrainment of a population of neurons (Herrmann, 2001) or due to linear superposition of SEPs in response to each pulse of the train (Capilla et al., 2011). Critically, both explanations could explain the increase in EEG/MEG power and inter-trial phase coherence (ITPC) at the frequency of stimulation (Capilla et al., 2011; Keitel et al., 2014). When refractoriness is taken into account there is compelling evidence for the superposition hypothesis (Capilla et al., 2011; Colon et al., 2012). Nevertheless, there is reason to believe both mechanisms coexist (Colon et al., 2012; Notbohm et al., 2016). Oscillating systems have a preferred frequency and when matched by the external stimulus train the system resonates (Colon et al., 2012; Herrmann, 2001; Notbohm et al., 2016; Vialatte et al., 2010). Therefore, at certain frequencies the signal recorded could be largely because of entrainment rather than rhythmic SEPs (Colon et al., 2012). Hence, in our study we delivered trains of MNS at 12 and 20Hz during concurrent MEG recording to investigate entrainment effects (adapted from (Morera Maiquez et al., 2020b)). Replication of this experiment in MEG is important to ensure validity of the findings and to probe the effects in the somatosensory cortex. In MEG the signal is less affected by the conductivity of the overlying tissue compared to EEG (Baillet, 2017; Cheyne, 2013), and therefore the models used for source localisation are simpler due to the lower level of spatial smearing (Muthukumaraswamy, 2014). Furthermore, compared to EEG, MEG has lower susceptibility to interference from non-neuronal sources such as muscles (Boto et al., 2019). Here we hypothesise that there will be entrainment of oscillations within the contralateral somatosensory cortex during rhythmic, but not arrhythmic, MNS. To conclude that rhythmic MNS induces entrainment we will need to show (i) a frequency and region-specific increase in instantaneous amplitude, (ii) synchronisation of the phase with the external source and (iii) that these effects are unlikely to be due to rhythmic evoked potentials.

## 2. Experimental Procedures

- The MATLAB code for MNS delivery is available on OSF https://osf.io/28u6b/
- As the MRI data used during analysis were not collected by the researchers involved, we do not have ethical approval to share these.
- MEG data can be made available on request if a formal data sharing agreement is in place.

### 2.1. Subjects

Twenty healthy participants were recruited for the study. Nineteen participants were right handed according to the Edinburgh Handedness Inventory (Oldfield, 1971). Participants gave informed consent and the experimental paradigm received local ethics committee approval (School of Psychology, University of Nottingham). Participants agreed that pre-existing structural MRI data (obtained within the Sir Peter Mansfield Imaging Centre, University of Nottingham) could be used by the researchers. One participant (female, 21 years old, right-handed) was excluded prior to analysis due to excessive movement during the MEG recording, leaving 19 usable datasets (aged 26.7 ± 3.6 years (mean ± SD); 11 female). A small inconvenience allowance was provided for volunteers for their participation.

### 2.2. Median Nerve Stimulation

Stimulation was delivered with electrodes (cathode proximal) positioned on the right forearm over the median nerve, using a Digitimer constant current stimulator model DS7A (Digitimer Ltd, UK). Pulse width was set at 0.2ms and maximum compliance voltage (Vmax) was 400V. Participants were seated and told to rest their forearm on the chair armrest to ensure the muscles were relaxed. The stimulation threshold was set at the minimum intensity required for a visible thumb twitch (mean ± SD) 9.1 ± 3.2 mA.

Four hundred trials of stimulation were delivered to the right median nerve during concurrent MEG recording, with a short break every 100 trials. Each trial consisted of 10 pulses delivered at the frequency of interest, rhythmic 12Hz and 20Hz in the test conditions. Arrhythmic controls were used to ensure that similar responses did not occur with random stimulation at the same average frequency. During the control trials the same number of pulses were delivered as in the test condition, but with a random inter-stimulus interval (ISI) which was on average the same as the rhythmic trials (but constrained to a minimum ISI of 0.01s). The arrhythmic patterns were not the same for all trials or participants, however, the first pulse was always delivered at time 0 and the last pulse was always delivered at the end of the train (450 or 750ms), regardless of the condition. For each of these 4 conditions, 100 trials were randomly delivered with an inter-trial interval of 4s using an in-house Matlab script (Matlab R2012a, Mathworks, Natick, MA) (Figure 1).

**Figure 1.**
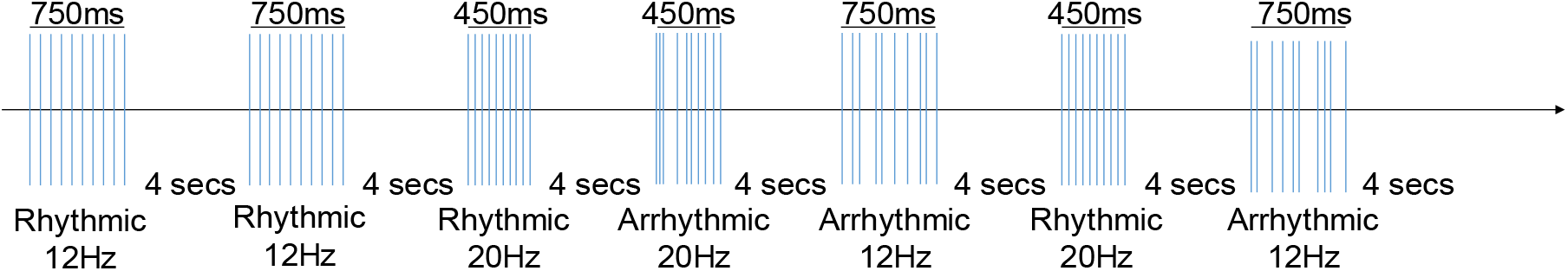
Stimulation overview. A diagram showing an example of the trial setup for the MNS.

### 2.3. EMG Measurement

Electromyography (EMG) electrodes were placed over the right abductor pollicis brevis (APB) muscle in a belly tendon montage to allow motor evoked potentials (MEPs) to be recorded for twenty pulses of MNS at the intensity for a minimum thumb twitch, with 3s between each pulse. A Brainamp ExG (Brain Products GmbH, Gilching, Germany) was used to amplify the signal and Brain Vision Recorder (Brain Products GmbH, Gilching, Germany) was used to record the EMG data (bandpass filtered 10-2000Hz, sampling rate 5kHz). Peak-to-peak amplitudes were measured using an in-house Matlab script to determine the baseline MEP amplitude (mean = 1219.13μV, range = 52.10 - 7490.9μV) (Matlab R2017a, Mathworks, Natick, MA). EMG data from 6 individuals was not recorded due to technical issues.

### 2.4. MEG Data Acquisition

MEG data were collected at the Sir Peter Mansfield Imaging Centre, University of Nottingham using a 275-channel CTF MEG system (MISL, Coquitlam, Canada) with participants in a seated position. The system was operated in a third order synthetic gradiometer configuration and data were sampled at 600Hz. Fiducial marker coils were placed on the nasion and bilateral preauricular points of the participants, to track head movement in relation to the MEG sensors. A Polhemus FASTRAK (Polhemus Inc, Vermont) was used to digitise the participants’ head shape and the relative positions of the fiducial markers. The data were examined by eye using commercial MEG data visualisation software (CTF MEG, Canada) and trials containing large artefacts were discarded, as were trials where the head moved more than 7mm from its initial position. One occipital channel was removed during preprocessing in one subject. Following preprocessing there were on average (mean ± SD) 7 ± 8 rhythmic 20Hz, 9 ± 10 arrhythmic 20Hz, 8 ± 9 rhythmic 12Hz and 8 ± 8 arrhythmic 12Hz trials removed per subject. The data were then segmented such that an epoch started 1s before stimulation and ended 3s following the end of stimulation. The epoch lengths of alpha and beta trials were 4.75s and 4.45s, respectively. An anatomical MRI scan (1.0mm^3^ resolution, MPRAGE sequence) from each subject was used for co-registration to the digitised head shape.

### 2.5. Data Analysis

After pre-processing, a linear-constraint minimum-variance (LCMV) beamformer was applied to the data (Van Veen et al., 1997). Using FLIRT (FMRIB’s Linear Image Registration Tool) (Jenkinson et al., 2012), a MNI (Montreal Neurological Institute) template brain was warped with respect to the subject’s downsampled (4mm) anatomical scan, as was the AAL (Automated Anatomical Labelling) atlas (Tzourio-Mazoyer et al., 2002). This allowed the cortex to be parcellated into 78 regions and the coordinates of the centroids from each region to be determined for each individual (Gong et al., 2009). Covariance was calculated for the entire experimental time window within a 1-150Hz frequency window (Brookes et al., 2008). The covariance matrix of the filtered data was regularised using the Tikhonov method, with the regularisation parameter set at 5% of the maximum singular value. The forward model was computed using dipole approximation and a multiple local spheres head model (Huang et al., 1999; Sarvas, 1987). Dipoles were rotated in the tangential plane to determine the orientation which yielded the maximum signal-to-noise ratio. Beamformer weights were calculated for the centroid of each brain region resulting in 78 timecourses (O’Neill et al., 2017), which were filtered between 1 and 48Hz.

### 2.6. Time Frequency Spectrograms

To visualise amplitude changes across the trial, time frequency spectrograms (TFSs) were calculated. Data were filtered into overlapping frequency bands spanning from 4 to 50Hz to visualise the changes across frequency bands in the left somatosensory cortex as defined by the AAL atlas. The analytic signal for the data within each band was determined using a Hilbert transform (HT), the absolute value of which gave the instantaneous amplitude of the MEG signal at each point in time. Timecourses for each frequency band were then averaged across trials, before being normalised using the average amplitude during the control window (0.24-0.99s for 12Hz trials and 0.54-0.99s for 20Hz trials). TFSs were averaged across subjects.

We also calculated the average amplitude during the period of stimulation to investigate the differences between rhythmic and arrhythmic without averaging out the effects of the arrhythmic condition due to the random pulse timings. First the data were filtered into the frequency band of stimulation (11-13Hz or 19-21Hz). Then the data from each trial was normalised using the average amplitude during the control window for that trial. Next, we calculated the mean amplitude within the period of stimulation and finally averaged across trials and subjects. The average amplitude from the periods of rhythmic and arrhythmic stimulation were then compared using a one-tailed Wilcoxon signed rank test.

### 2.7. Inter-Trial Phase Coherence

The timeseries from the contralateral somatosensory cortex were filtered into the same frequency bands as were previously used for the TFSs calculation. Following a HT of the data, the ITPC was calculated using formula (1) below:

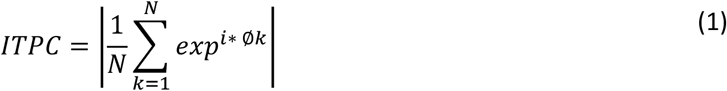

Where *N* is the number of trials and *Φk* is the phase angle in radians of the datapoint in the current trial.

### 2.8. Sensory Evoked Potentials

Rhythmic sensory evoked potentials would lead to an increase in instantaneous amplitude and ITPC during rhythmic stimulation, providing an alternative mechanism to entrainment (Thut et al., 2011b). To investigate whether this was the case, we looked at whether there was a full oscillation at the frequency of stimulation associated with each pulse of the rhythmic stimulation train to determine whether entrainment had occurred. To do this the beamformed signal was filtered between 8-12Hz for the 12Hz stimulation trials and 13-30Hz for the 20Hz stimulation trials. The data following the pulse timings (83.3ms for the 12Hz trials and 50ms for the 20Hz trials) were averaged for each of the 10 pulses across all trials for every subject, before being averaged across subjects. The same method was used to look at the broadband (1-48Hz) signal following each pulse, to investigate whether there was a SEP associated with each MNS pulse. Data from 1 participant were not included in the 20Hz condition analysis due to a loss of data, meaning we were unable to determine which trials were deleted during preprocessing, and as such could not determine the pulse timings for the remaining trials.

### 2.9. Statistical Analysis

For both the instantaneous amplitude and ITPC, timecourses were filtered according to the frequency of stimulation (11-13Hz or 19-21Hz) and were statistically compared using a one-tailed Wilcoxon signed rank test at each timepoint. Multiple comparisons were corrected for using the false discovery rate (FDR) of 0.05 (Benjamini and Hochberg, 1995; Benjamini and Yekutieli, 2001; Groppe, 2020). The same method was used to compare the timecourses following each pulse of rhythmic and arrhythmic stimulation for the evoked potential analysis. Post-hoc timecourse analysis was completed in the same manner but with a two-tailed test. All timecourse plots show the standard error of the mean (SEM) (Martínez-Cagigal, 2020).

## 3. Results

All analyses were conducted on the contralateral (left) *somatosensory* cortex as defined by the AAL atlas. Results for similar analyses on the contralateral *motor* cortex can be found in the supplementary information.

### 3.1. Entrainment: Increase in Instantaneous Amplitude

During the delivery of 10 rhythmic pulses at 12Hz, there was an increase in oscillatory amplitude at 12Hz (Figure 2A) within the contralateral sensorimotor cortex. This increase was not evident during arrhythmic stimulation where a decrease in amplitude was seen bilaterally in the sensorimotor cortices (Figure 2B). Similarly, during the 20Hz rhythmic condition there was an increase in the oscillatory amplitude at 20Hz (Figure 2C) within the contralateral sensorimotor cortex, which was not evident during arrhythmic stimulation at the same average frequency (Figure 2D). The ipsilateral sensorimotor cortex appeared to show a larger decrease in amplitude than the contralateral cortex during arrhythmic stimulation. This suggests that although there is the bilateral event related desynchronisation which is to be expected with somatosensory stimulation, the arrhythmic stimulation is lessening the extent of this in the contralateral cortex.

**Figure 2.**
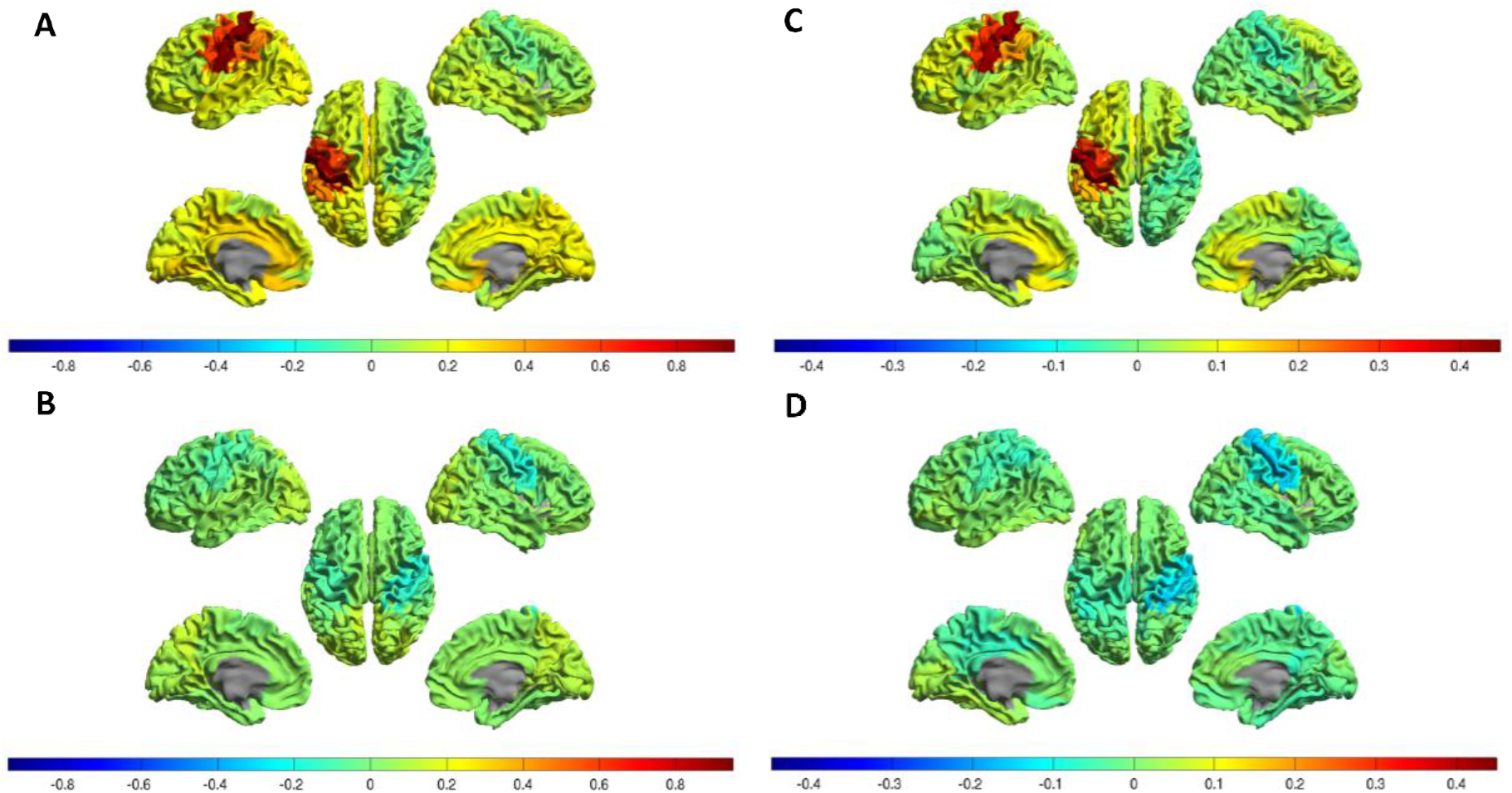
Localisation of the source of oscillatory changes at the frequency of interest compared to baseline. Oscillatory amplitude for 78 cortical regions of the AAL atlas during A) rhythmic 12Hz MNS and B) arrhythmic 12Hz MNS, C) rhythmic 20Hz MNS and D) arrhythmic 20Hz MNS.

Evidence of rhythmic entrainment comes from the TFSs where there was an initial broadband increase in amplitude associated with the first pulse (1s) of both rhythmic (Figure 3A, Figure 3D) and arrhythmic stimulation (Figure 3B, Figure 3E), which became specific to a narrowband around the frequency of stimulation during only rhythmic stimulation (Figure 3A, Figure 3D). This increase in amplitude returned to baseline shortly after the last pulse in the rhythmic condition. When the arrhythmic TFS is subtracted from the rhythmic TFS, it is evident that the difference between trials occurred only during the delivery of stimulation (Figure 3C, Figure 3F). The 12Hz relative amplitude within the window of stimulation was significantly higher in the rhythmic condition compared to arrhythmic (p≤0.05 between 1093 and 1883ms, FDR corrected). Similarly, the 20Hz relative amplitude significantly increases within the window of stimulation in the rhythmic condition compared to arrhythmic (p≤0.05 between 1127 and 1557ms, FDR corrected). As the arrhythmic condition involved pulses being delivered at different times during every trial, amplitude changes may have been averaged out during calculation of the TFS. Therefore, we calculated the average amplitude at the frequency of stimulation during the period that pulses were delivered for each trial before averaging across trials. This showed a significantly higher 20Hz amplitude in the rhythmic compared to the arrhythmic condition (t=182, z=3.48, p≤0.001). Analysis of the average amplitude across trials in the 12Hz condition again showed significantly higher 12Hz amplitude in the rhythmic condition (t=189, z=3.76, p≤0.001).

**Figure 3.**
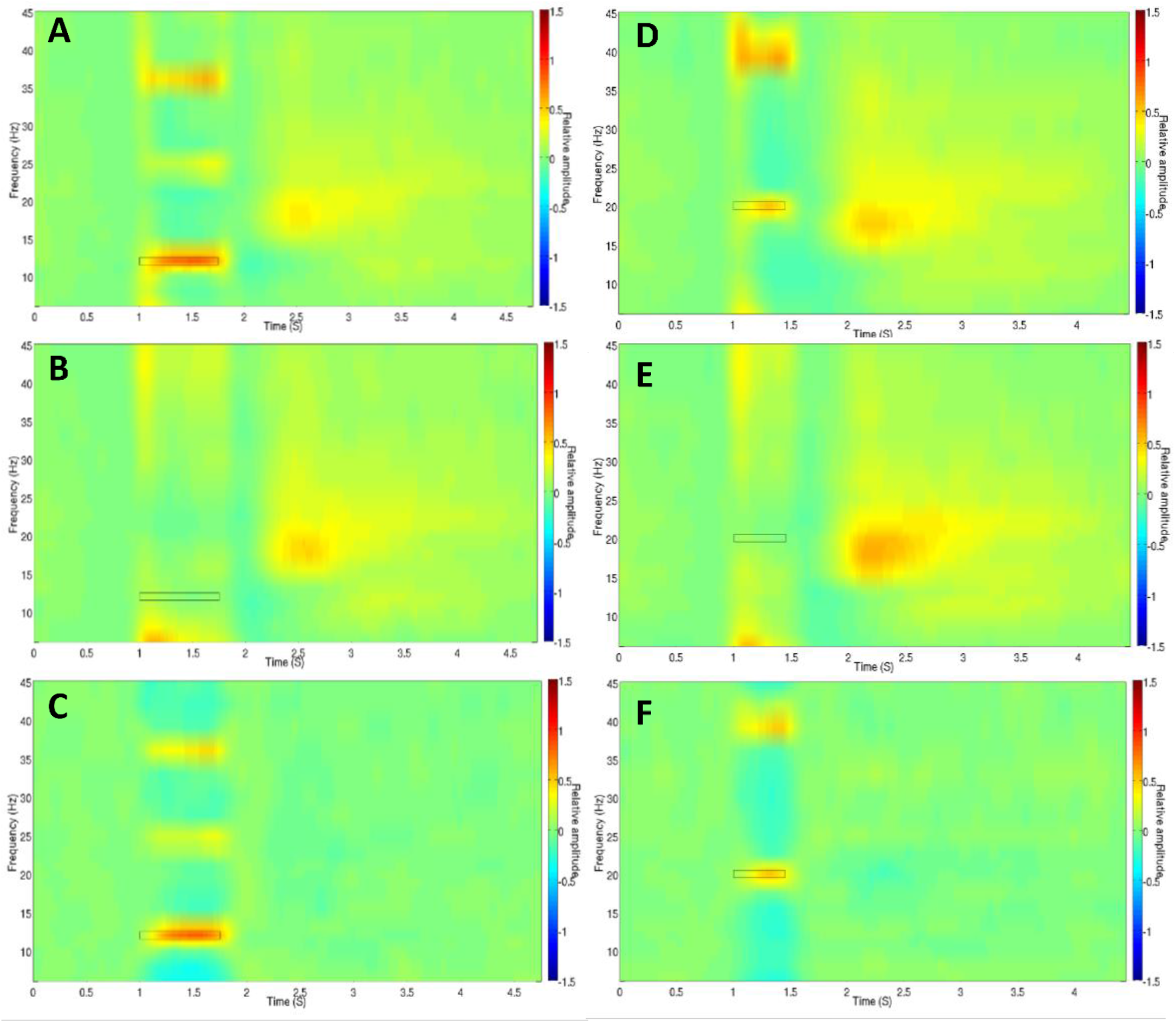
Figure 3. Relative amplitude in the contralateral somatosensory cortex. (A-C) TFSs of relative amplitude changes when 12Hz MNS was delivered between 1 and 1.75s in either a A) rhythmic or B) arrhythmic pattern with C) showing the difference in amplitude between the two conditions. (D-F) TFSs of relative amplitude changes when 20Hz MNS was delivered between 1 and 1.45s in a D) rhythmic E) arrhythmic pattern with F) showing the difference in amplitude between the two conditions.

### 3.2. Entrainment: Increase in Phase Coherence

For both rhythmic and arrhythmic stimulation, there was an initial increase in ITPC across a wide range of frequencies (Figure 4). After the first stimulation pulse, this increase in ITPC was maintained in only the rhythmic condition and centred on the frequency of stimulation and its harmonics (Figure 4A, Figure 4C). The ITPC within the window of stimulation was significantly higher in the rhythmic condition compared to arrhythmic (12Hz: p≤0.05 between 1047-1990ms, FDR corrected) (20Hz: p<0.05 between 960-1700ms and at 1703ms, FDR corrected).

**Figure 4.**
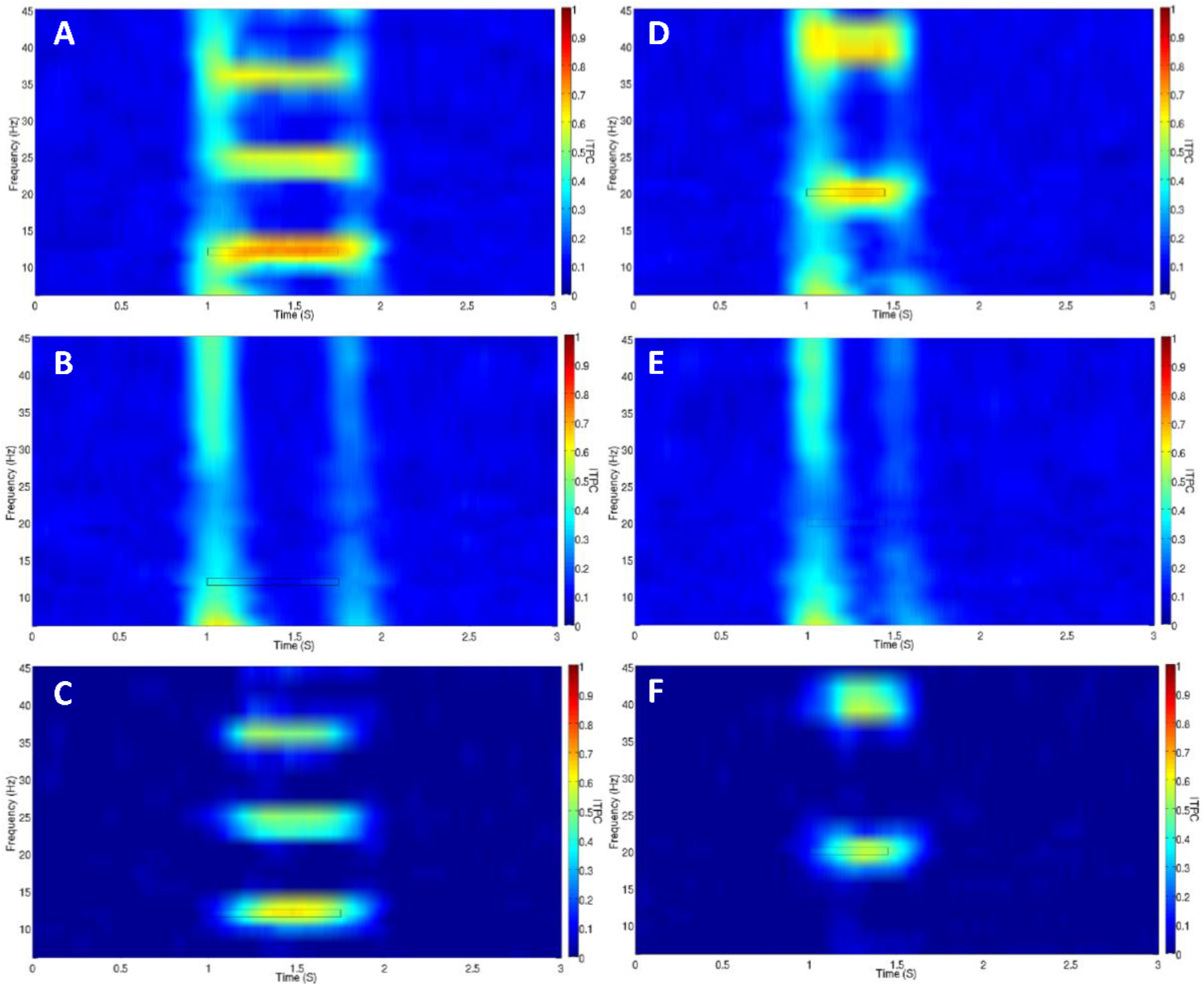
Inter-trial phase coherence in the contralateral somatosensory cortex. (A-C) TFSs of ITPC when 12Hz MNS was delivered between 1 and 1.75s in A) rhythmic B) arrhythmic pattern with C) showing the difference in ITPC between the two conditions. (D-F) TFSs of ITPC when 20Hz MNS was delivered between 1 and 1.45s in a D) rhythmic E) arrhythmic pattern with F) showing the difference in ITPC between the two conditions.

### 3.3. Entrainment: Hemispheric Specificity

To investigate whether the effect was hemisphere specific, as suggested by the results shown in Figure 2, we analysed the relative amplitude at the frequency of stimulation in the right and left somatosensory cortices during the rhythmic trials. The increase in relative amplitude during the 12Hz rhythmic stimulation (p≤0.05 between 1105 and 1842ms, FDR corrected) and the subsequent rebound (p≤0.05 for 3715-3960ms, 4397-4405ms and at 4417ms, FDR corrected) was significantly higher in the left somatosensory cortex, which is contralateral to the stimulated arm. Similarly, the increase in amplitude in the left hemisphere during the 20Hz rhythmic stimulation was significantly higher (p≤0.05 between 1152-1520ms, FDR corrected), as was the subsequent rebound (p≤0.05 for 1993-3165ms, 3168-3173ms, 3223-3373ms, 3632-3637ms, 3643-3867ms, 3875-3902ms and 3968-4092ms, FDR corrected).

### 3.4. Sensory Evoked Potentials

A possible explanation for the increase in instantaneous amplitude and ITPC seen during stimulation is that there was a SEP associated with each pulse of the MNS (Thut et al., 2011b). Therefore, we investigated whether each rhythmic pulse was associated with a full oscillation at the frequency of stimulation and whether this could be explained by the filtering of an SEP (Figure 5). Each subplot contains the data following the pulse timings in each condition for a complete oscillation at the frequency of stimulation meaning a timeframe of 83.3ms was used for the 12Hz trials and 50ms for the 20Hz trials. As is evident in the 12Hz trials a full oscillatory cycle was associated with each pulse of both the rhythmic and arrhythmic trains. However, on investigation of the broadband 1-48Hz data in the 12Hz trials, an evoked component was clearly associated with each pulse (Figure 5B). Following the SEP evoked by the first pulse, the remainder of the evoked potentials appear to be smaller in amplitude. However, with the exception of the initial MNS pulse, the pulses of the 20Hz trials do not appear to be associated with a SEP (Figure 5D). In Figure 5A and Figure 5C the timings were shifted by 20ms to account for the approximate time it would take for the afferent volley to reach the cortex.

**Figure 5.**
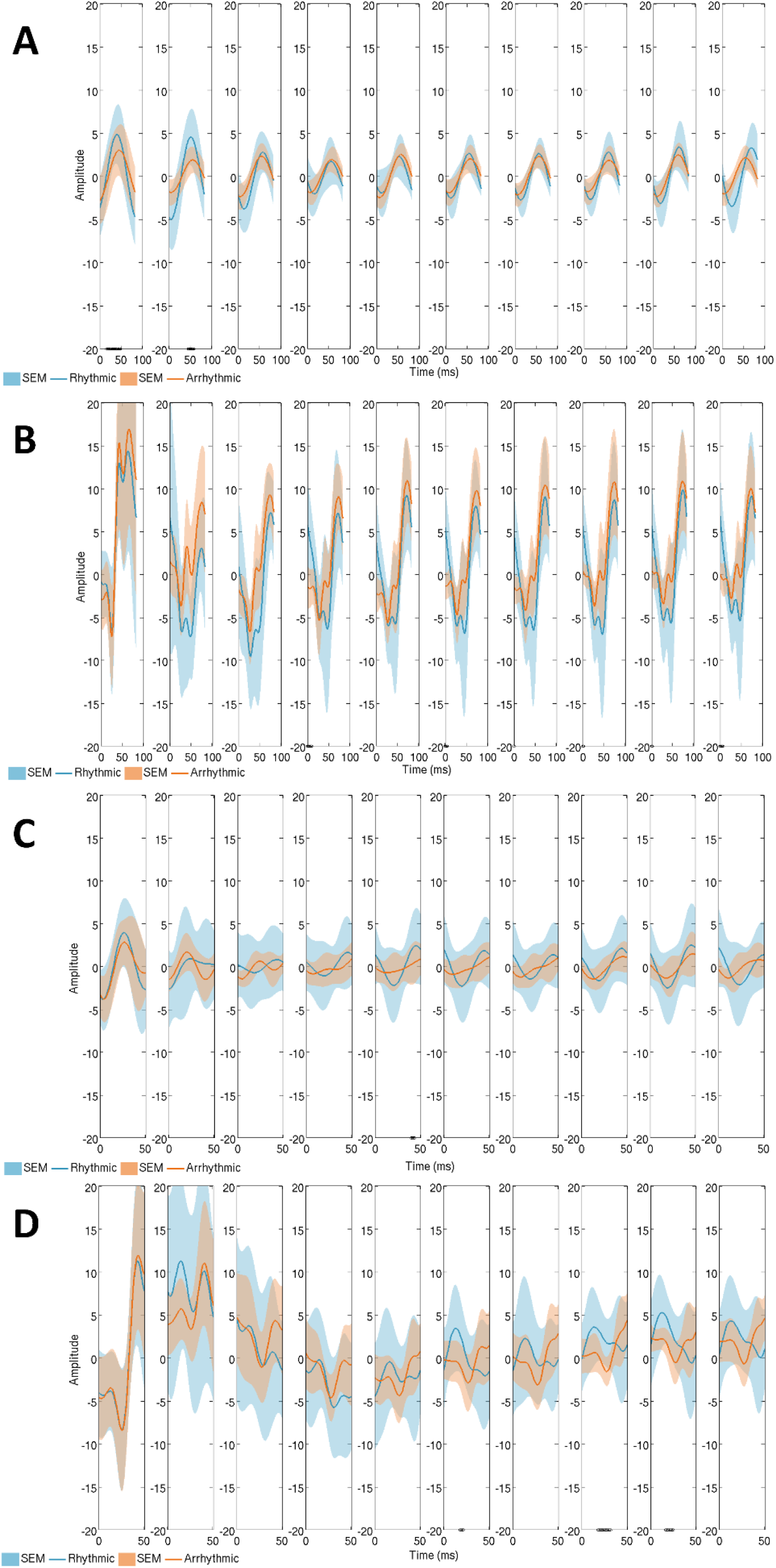
Evoked components in the contralateral somatosensory cortex. A figure showing: A) The 8-12Hz oscillatory response to each pulse of the rhythmic and arrhythmic 12Hz MNS. B) An evoked component associated with each pulse of the rhythmic and arrhythmic 12Hz MNS. C) The 13-30Hz oscillatory response to each pulse of the rhythmic and arrhythmic 20Hz MNS. D) An evoked component associated with the first pulse of the rhythmic and arrhythmic 20Hz MNS and the progressive entrainment of beta oscillations in the rhythmic condition. A black line along the x-axis marks timepoints where a significant difference (p≤0.05) is seen (FDR corrected).

The data in Figure 5B and Figure 5C were not shifted and the negative deflection at ~20ms (N20 peak) reflects when the signal arrived in the primary somatosensory cortex (Passmore et al., 2014).

### 3.5. Post-hoc Analyses

Due to the evidence of aftereffects seen in the paper by Morera and colleagues (Morera Maiquez et al., 2020b), we also investigated the broadband mu-alpha (8-12Hz) and beta (13-30Hz) effects following both 12Hz and 20Hz stimulation. The only significant effect was in the mu-alpha (8-12Hz) band during 20Hz arrhythmic stimulation compared to rhythmic stimulation, indicating a greater degree of suppression in the rhythmic trials as seen in Figure 3(D-F) (p≤0.05 for 1123-1353ms and 1387-1487ms, FDR corrected). There were no significant differences in the aftereffects of rhythmic and arrhythmic stimulation.

As the beneficial effects of rhythmic MNS on tics is not restricted to the stimulated limb, as demonstrated in a recent paper by Morera and colleagues, we decided to investigate the spread of the effect using a voxel LCMV beamformer (Morera Maiquez et al., 2020b). This revealed that the focus of the increase in amplitude was in the contralateral sensorimotor hand region (Supplementary information).

## 4. Discussion

We investigated whether rhythmic MNS could be used to entrain oscillations at frequencies associated with sensorimotor inhibition, mu-alpha (12Hz) and beta (20Hz). Our results demonstrate that rhythmic 12Hz stimulation resulted in an increase in the relative amplitude and ITPC at 12Hz for the duration of the stimulation in the contralateral somatosensory cortex. Both of these findings can be split into two segments: 1) an increase in both amplitude and ITPC across a broad range of frequencies for the first pulse in both the rhythmic and arrhythmic trials; 2) an increase specific to the frequency of stimulation for the remainder of the pulse train, which was only seen in the rhythmic condition. However, we also show that for both rhythmic and arrhythmic 12Hz MNS, an evoked component was associated with each pulse of the stimulation. The effects on relative amplitude and ITPC were replicated in the 20Hz condition, however the presence of evoked potentials was less clear. These results, and the evidence they provide as to whether MNS can entrain neuronal oscillations, are discussed below.

### 4.1. Steady-State Evoked Response

Thut et al., 2011 previously showed that ~10Hz TMS caused an initial broadband increase in amplitude in both the rhythmic and arrhythmic stimulation conditions, before progressive entrainment was seen in the rhythmic condition (Thut et al., 2011b). The authors proposed that the delivery of the first few pulses may have resulted in phase resetting of neural generators within the cortex whose frequencies of oscillation varied across a broad range of frequency bands (Thut et al., 2011b). Therefore, phase resetting of these oscillators could have resulted in the initial broadband increase in amplitude. The subsequent pulses however will have only occurred in phase with generators oscillating in the alpha band leading to progressive synchronisation of oscillations at this frequency (Thut et al., 2011b).

Here we show that MNS causes an initial broadband increase in relative amplitude, suggesting that peripheral stimulation triggers an initial phase reset of the neural generators within the contralateral somatosensory cortex regardless of the pattern of stimulation. In line with the theory of entrainment we see that the remainder of the rhythmic pulses cause a frequency specific increase in amplitude. This is also mirrored in the ITPC results, where the initial broadband increase aligns with the theory that numerous neural generators phase reset resulting in phase coherence across trials. As the remaining rhythmic pulses are associated with an increased ITPC in the stimulated frequency band (and its harmonics) this shows that oscillations across trials are relatively more in phase during the stimulation suggesting synchronisation with the pulses of MNS.

However, one central issue in concluding that a rhythmic oscillatory response to a rhythmic stimulus is neural entrainment is whether the same data could be explained by rhythmic evoked potentials (For in-depth review please see (Thut et al., 2011a; Zoefel et al., 2018)). If we deliver a stimulus at 12Hz it is expected that we would see a response 12 times a second (Zoefel et al., 2018). An investigation of this possibility by Capilla and colleagues showed that the oscillatory response seen during rhythmic stimulus presentation can be modelled through linear superposition of evoked potentials, which casts doubt on the explanation of entrainment (Capilla et al., 2011). Therefore, rhythmic SEPs mask at least in part any entrainment effect rhythmic MNS has on endogenous oscillations in the 12Hz condition. As both responses are expected to repeat at the same frequency, this makes it difficult to determine whether there is underlying entrainment (Zoefel et al., 2018). Furthermore, we would expect oscillations to continue after stimulation if entrainment had taken place (Thut et al., 2011a; Zoefel et al., 2018). During entrainment oscillators become phase-aligned to the external source and, as phase is a free parameter, any change in phase alignment should only occur if the system is perturbed (Pikovsky et al., 2003). Here, we see a short-lived continuation of the increase in amplitude after the last pulse of MNS, but this could be due to the evoked potential associated with this last pulse.

In contrast to the 12Hz condition, the 20Hz rhythmic stimulation appears to show oscillations within the broadband 1-48Hz data following the evoked potential associated with the first MNS pulse, suggesting that entrainment may have occurred. Previous experiments investigating steady-state responses of the somatosensory cortex have reported a maximal amplitude between 21 and 27 Hz (Müller et al., 2001; Snyder, 1992; Tobimatsu et al., 2000, 1999). In recent preprint, Walti and colleagues reported that somatosensory oscillations could be entrained by vibrotactile stimulation at beta but not alpha frequency (Wälti et al., 2019 (preprint)). They concluded that their results could be explained by beta being the preferred frequency of the somatosensory system, a theory that our results also support.

Nevertheless, the evoked potentials and entrained oscillations are generated by synchronous firing of neuronal populations within the contralateral somatosensory cortex meaning these neurons are ‘occupied’ in processing of the afferent input. Recent research has demonstrated the possibility that rhythmic MNS could be therapeutically beneficial in reducing tic frequency in TS patients (Morera Maiquez et al., 2020b). It is thought that there is a high level of ‘sensorimotor noise’ associated with the occurrence of tics in TS leading to a difficulty in discriminating between the signals preceding voluntary and involuntary movement (Ganos et al., 2015). Increasing synchronised firing of neuronal populations within the sensorimotor cortex through MNS may lead to a decrease in this noise. Alternatively, it is known that ~20ms following median nerve stimulation there is a decrease in the amplitude of MEPs elicited by a TMS pulse to the contralateral motor cortex in a process known as short afferent inhibition (SAI) (Tokimura et al., 2000). As there is thought to be a deficit in this form of inhibition in TS patients it could be that continuous MNS compensates for this deficit and aids in the inhibition of tics and urges (Morera Maiquez et al., 2020b; Orth et al., 2005; Orth and Rothwell, 2009). Considering that the research into the therapeutic potential of MNS used stimulation at 10Hz, it is interesting that evoked potentials were elicited in the sensory cortex with both rhythmic and arrhythmic 12Hz stimulation (Morera Maiquez et al., 2020b). It is therefore of interest as to whether arrhythmic stimulation also reduces the frequency of tics i.e. is the beneficial effect due to SAI and/or a decrease in sensorimotor noise, or is the rhythmicity of the stimulation important (Morera Maiquez et al., 2020b).

### 4.2. Aftereffects of Rhythmic Stimulation

The observance of a desynchronisation and rebound of sensorimotor oscillations following MNS trains is typical of that seen following one pulse of MNS (Pfurtscheller, 1981). The only significant difference in the effect seen here was greater desynchronization in the 8-12Hz band during 20Hz rhythmic stimulation compared to arrhythmic. We found no difference in aftereffects. This contrasts with recent findings which indicated that there was greater mu-alpha and beta desynchronization and increased beta rebound following 19Hz arrhythmic stimulation (Morera Maiquez et al., 2020a (preprint)). On inspection of the data reported by Morera and colleagues, there is a short period of increased mu-alpha suppression towards the end of the stimulus train and for a short time following rhythmic stimulation, similar to what is seen in our data. This suggests that the typical mu-alpha band desynchronization seen during movement is maintained in the rhythmic condition. However, further findings by the same authors also describe greater beta desynchronization and increased beta rebound following 12Hz arrhythmic stimulation (Morera Maiquez et al., 2020b). Our experiment along with the two previous studies left only 4 seconds between each trial during which the post-movement beta rebound may still be ongoing (Pakenham et al., 2020), critically this is also the period during which the experiments defined their control windows. This means that the baseline was likely contaminated by higher beta amplitude, the extent to which likely depended on both the sample group and the condition of the preceding trial, which in the current study could have been either 12 or 20Hz. Investigation of the uncorrected TFS confirms a high beta amplitude continues throughout the control window. However, given that the intertrial interval of 4 seconds was the same in both the previous and the current study, and the same for both rhythmic and arrhythmic trials, then any differences are unlikely to be explained by the inter-trial interval alone (Morera Maiquez et al., 2020b). Another potential factor is the subjective nature of the visible thumb twitch which may have led to participants in one study to experience higher intensities of stimulation. Ultimately further investigation into the aftereffects of rhythmic versus arrhythmic MNS is required. Future studies should ideally use longer intertrial intervals to allow for beta rebound (Pakenham et al., 2020), and if possible an objective method of thresholding, for example, through measurement of MEPs as is common practice with TMS.

### 4.3. Conclusion

To conclude, the evidence from this research suggests that endogenous oscillations within the somatosensory cortex can be entrained by 20Hz rhythmic MNS. On the other hand, the oscillatory effects of 12Hz rhythmic MNS are largely caused by the presence of rhythmic SEPs. As SEPs are present in both the rhythmic and arrhythmic 12Hz conditions, investigation of the effect of arrhythmic stimulation on tics and urges is warranted. Nonetheless, the behavioural effects of rhythmic peripheral nerve stimulation are clinically important, however a better understanding of how these behavioural effects are produced is vital for fine-tuning their development as a therapeutic technique (Morera Maiquez et al., 2020b).

## Abbreviations

AAL: Automated Anatomical Labelling
APB: abductor pollicis brevis
EMG: electromyography
FDR: false discovery rate
FLIRT: FMRIB’s Linear Image Registration Tool
HT: Hilbert transform
ISI: inter-stimulus interval
ITPC: inter-trial phase coherence
LCMV: linear-constraint minimum-variance
MEPs: motor evoked potential
MNI: Montreal Neurological Institute
MNS: median nerve stimulation
NIBS: non-invasive brain stimulation
SAI: short afferent inhibition
SEP: sensory evoked potential
SMA: supplementary motor area
SSEP: steady-state evoked potential
tACS: transcranial alternating current stimulation
TFS: time frequency spectrogram
TMS: transcranial magnetic stimulation
TS: Tourette’s syndrome

## Author Contributions

**MH.** Conceptualisation, Experimental design, Data analysis, Data interpretation, Writing paper - Original Draft.

**BM.** Conceptualisation, Experimental design, Data interpretation, Writing paper – Review and Editing.

**MB.** Data analysis, Data interpretation, Writing paper – Review and Editing.

**SJ.** Conceptualisation, Experimental design, Data interpretation, Writing paper – Review and Editing.

## Acknowledgments

Data collection for this paper was supported by research grant from NIHR Nottingham Biomedical Research Centre. The views expressed are those of the authors and not necessarily those of the NHS, the NIHR or the Department of Health.

