## Supplementary Information for "The Oscillatory Effects of Rhythmic Median Nerve Stimulation"

Analysis of amplitude (Figure S.1) and ITPC (Figure S.2) within the contralateral motor cortex shows similar results to those seen for the sensory cortex in our main manuscript, with an increase during the rhythmic pattern of stimulation for both the 12 and 20Hz conditions.

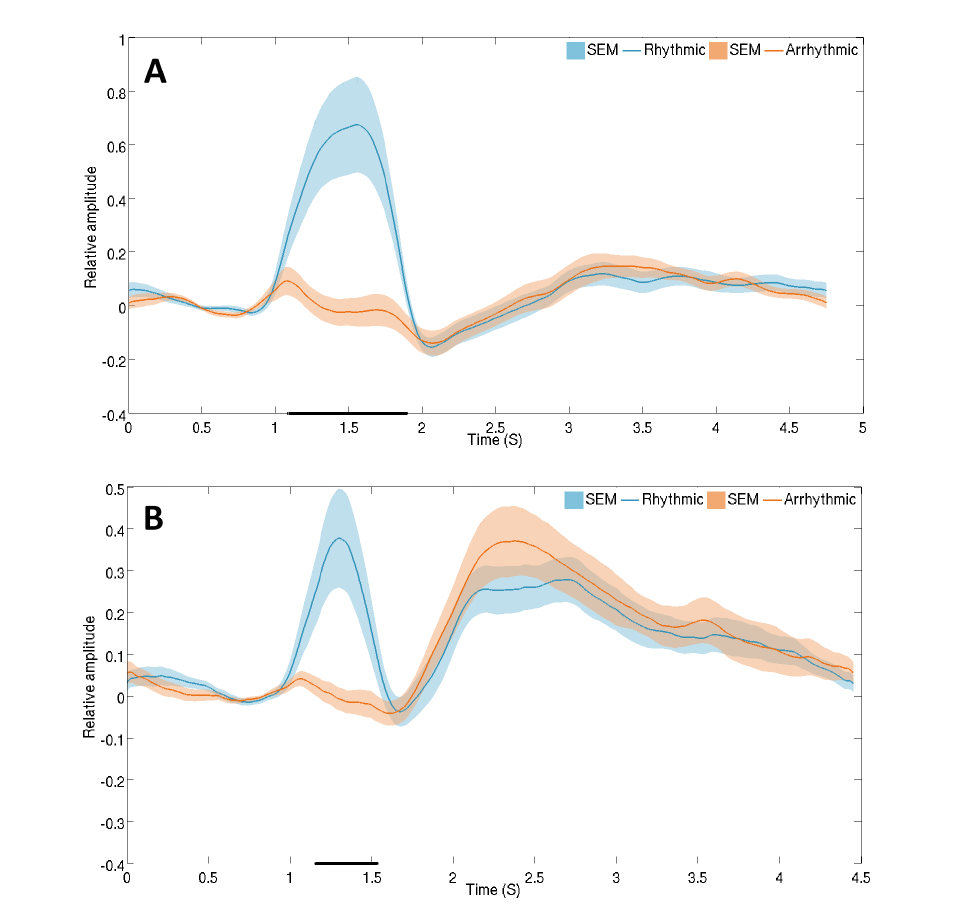

**Figure S.1. Comparison between rhythmic and arrhythmic amplitude changes in the contralateral motor cortex.**

Graphs showing the difference in the relative A) 12Hz amplitude and B) 20Hz amplitude in the rhythmic and arrhythmic conditions. A black line along the x-axis marks timepoints where a significant difference (p≤0.05) is seen (FDR corrected).

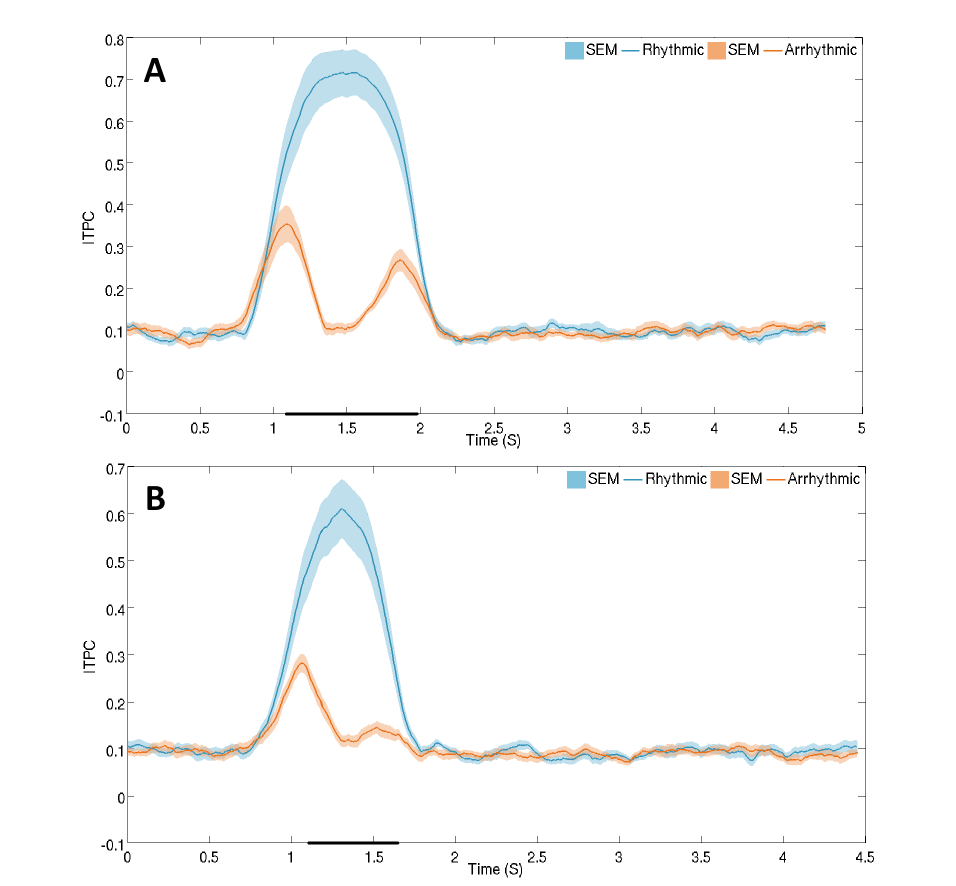

**Figure S.2. Inter-trial phase coherence in the contralateral motor cortex**.

A graph showing the difference between ITPC in the rhythmic and arrhythmic conditions during A) 12Hz and B) 20Hz stimulation. A black line along the x-axis marks timepoints where a significant difference (p≤0.05) is seen (FDR corrected).

**Focus in contralateral hand region**

A LCMV beamformer was applied to the preprocessed data to allow comparison of the data in the active (20Hz [1s 1.45s]; 12Hz [1s 1.75s]) and passive (20Hz [0.54s 0.99s]; 12Hz [0.24s 0.99s]) windows, for rhythmic MNS at the frequency of stimulation (Van Veen et al., 1997). Beamformer weights were calculated for each voxel, to create a pseudo-T-statistic map for each condition. Covariance was calculated for the entire experimental time window in a frequency window set according to the frequency of stimulation (19-21Hz and 11-13Hz) (Brookes et al., 2008). The covariance matrix of the filtered data was regularised using the Tikhonov method, with the regularisation parameter set at 4% of the maximum singular value. Individual maps were sampled on a 4mm grid of the subject’s anatomical scan. For the group average (Figure S.3), individual maps were subsequently transformed to MNI coordinate space using FLIRT. The neural generator of the entrained oscillations was identified using the peak amplitude in the frequency band of interest on the pseudo-T-statistic map. This revealed that the focus of the amplitude increase during rhythmic stimulation was in the contralateral sensorimotor hand region.

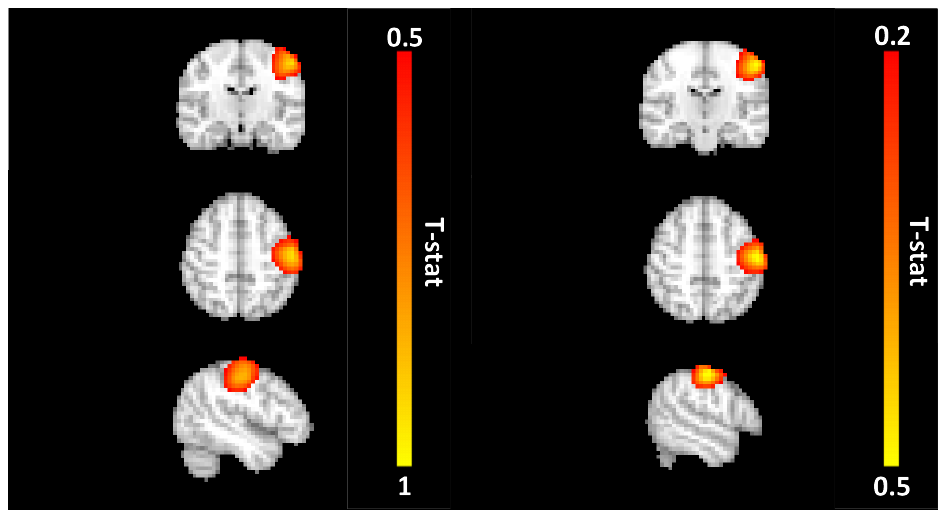

**Figure S.3. Pseudo-t-statistical map**

A figure showing the average location of the amplitude increase during rhythmic stimulation at 12 (left) and 20Hz (right) localises to the contralateral sensorimotor hand region (MNI space) (radiological orientation).
